# Primary cilia in osteoblasts and osteocytes are required for skeletal development and mechanotransduction

**DOI:** 10.1101/2023.12.15.570609

**Authors:** Mariana Moraes de Lima Perini, Julie N Pugh, Elizabeth M Scott, Karan Bhula, Austin Chirgwin, Olivia N Reul, Nicolas F Berbari, Jiliang Li

## Abstract

Primary cilia have been involved in the development and mechanosensation of various tissue types, including bone. In this study, we explored the mechanosensory role of primary cilia in bone growth and adaptation by examining two cilia specific genes, IFT88 and MKS5, required for proper cilia assembly and function. To analyze the role of primary cilia in osteoblasts, Osx1-GFP:Cre mice were bred with IFT88^LoxP/LoxP^ to generate mice with a conditional knockout of primary cilia in osteoblasts. A significant decrease in body weight was observed in both male (p=0.0048) and female (p=0.0374) conditional knockout (cKO) mice compared to the wild type (WT) controls. The femurs of cKO mice were significantly shorter than that of the WT mice of both male (p=0.0003) and female (p=0.0019) groups. Histological analysis revealed a significant difference in MAR (p=0.0005) and BFR/BS (p<0.0001) between female cKO and WT mice. The BFR/BS of male cKO mice was 58.03% lower compared to WT mice. To further investigate the role of primary cilia in osteocytes, Dmp1-8kb-Cre mice were crossed with MKS5^LoxP/LoxP^ to generate mice with defective cilia in osteocytes. In vivo axial ulnar loading was performed on 16-week-old mice for 3 consecutive days. The right ulnae were loaded for 120 cycles/day at a frequency of 2Hz with a peak force of 2.9N for female mice and 3.2N for male mice. Load-induced bone formation was measured using histomorphometry. The relative values of MS/BS, MAR and BFR/BS (loaded ulnae minus nonloaded ulnae) in male MKS5 cKO mice were decreased by 24.88%, 46.27% and 48.24%, respectively, compared to the controls. In the female groups, the rMS/BS was 52.5% lower, the rMAR was 27.58% lower, and the rBFR/BS was 41.54% lower in MKS5 cKO mice than the WT group. Histological analysis indicated that MKS5 cKO mice showed significantly decreased response to mechanical loading compared to the controls. Taken together, these data highlight a critical role of primary cilia in bone development and mechanotransduction, suggesting that the presence of primary cilia in osteoblasts play an important role in skeletal development, and primary cilia in osteocytes mediate mechanically induced bone formation.

## 2. Introduction

The primary cilium is a single, immotile, microtubule-based organelle present on nearly every cell in the body. It was first identified in the skeleton on chondrocytes over 50 years ago [1]. While it had been thought to be nothing but vestigial for its lack of motility, the primary cilium has recently come to the forefront of signaling studies and is now believed to be an essential regulator in many signaling pathways [2–5]. The cilium has shown to be indispensable for development, and plays a major role in mechanosensing in various tissue types, including the liver, kidney, endothelium, and bone [3].

Studies have shown that primary cilia possess properties that make them a great candidate for the coordination of signal transduction. These properties include receptors, ion channels, effector proteins, and transcription factors that can pass on stimuli from the extracellular environment to control development and homeostasis [5, 6]. For this reason, abnormal function or defects of the primary cilium have been linked to serious pathologies resulting in multisystemic defects, highlighting the importance of these organelles [4, 6].

The primary cilium has been reported to play a major role in the regulation of important pathways, such as Hedgehog (HH), Wnt, TGF-β, BMP, and Notch, all of which orchestrate skeletogenesis [2, 3, 7, 8]. With its involvement in so many pathways that are indispensable for tissue function and homeostasis, it is not a surprise that the primary cilium has been a focal research point across various medical fields.

The assembly of the primary cilium relies on the transport of axonemal precursors from the cell body to the ciliary tip [6]. Even though many proteins reside in the primary cilia, these ciliary proteins are not synthesized there and require a transport system [5]. This transport is referred to as intraflagellar transport (IFT). IFT is responsible not only for assembling the primary cilium but also for its maintenance. IFT is separated into two complexes, A and B, and with the help of motor proteins, it is responsible for transporting ciliary proteins to and from the cell. Motor proteins of the Kinesin-II family (KIF3A, KIF3B, and KAP3) transports proteins in the anterograde direction (base to tip), regulated by complex B consisting of about 14 IFT proteins, including IFT88. The retrograde (tip to base) transport is regulated by complex A, composed of six IFT proteins and dynein motor proteins (DYNC2H1 and DYNC2L1) [5, 6]. Between the basal body of the cell and the cilium is a zone referred to as the transition zone (TZ). This zone contains proteins that control the entrance and exit of proteins to and from the cilium [14, 15].

Intraflagellar transport 88 (Ift88), previously known as Tg737 or Polaris, is a gene that encodes for the IFT88 protein and is a key component of the IFT-B complex on primary cilia. Mutations disrupting any of the IFT components, including Ift88, impairs the formation of the primary cilium, most likely due to the inability of ciliary proteins to enter the primary cilium through the anterograde complex [16, 17].

Several protein complexes that localize in the TZ have been identified, including the Meckel Syndrome (MKS) and Nephronophthisis (NPHP) complexes. Mutations involving proteins both in IFT complexes as well as in the TZ can lead to absent or dysfunctional cilia by disrupting ciliary membrane composition, including the mislocalization of signaling proteins and loss of other TZ proteins [14, 15, 18, 19]. Located in the primary cilium’s TZ, retinitis pigmentosa GTPase regulator-interacting protein 1 like (RPGRIP1L), also referred to as MKS5, Ftm, and Nphp8, is a crucial gene for the function of the primary cilium. This protein may be able to regulate what is allowed in and out of the primary cilium and serve as an anchor between the cilium and the basal body [18, 20]. It is believed MKS5 holds a key role in the TZ, acting as a “bridge” between the MKS and NPHP modules. It is also believed that MKS5 may act as an “assembly factor” for all other MKS proteins and/or as an assembly factor for building the TZ [14, 18].

Increasing evidence suggests that the primary cilium may be involved in bone cell homeostasis, the bone’s response to mechanical loading, and bone mechanotransduction. Understanding he function and mechanism of the primary cilium in bone can pave the way for a new course of treatment for diseases associated with bone loss, ciliopathies, and fracture healing, making the primary cilium a new target organelle to treat these pathologies. In this study, we focused on two genes, IFT88 and MKS5, in two mouse models to investigate the role of the primary cilium in bone growth and mechanotransduction.

## 3. Materials and Methods

### 3.1 Animal Models

All animal procedures were reviewed and approved by the Institutional Animal Care and Use Committee (IACUC) at the Indiana University-Purdue University, Indianapolis (IUPUI).

Pre-osteoblast-specific deletion of the IFT88 gene was achieved with the breeding of Osx1-GFP::Cre (Stock No. 006361, Jackson Laboratories, Bar Harbor, ME, USA) positive mice and mice with LoxP sites flanking the IFT88 exon (Ift88^LoxP/LoxP^, Stock No. 022409, Jackson Laboratories, Bar Harbor, ME, USA) [21], generating Osx-Cre; IFT88^LoxP/LoxP^ (cKO mice), and Osx-Cre; IFT88^+/+^ (littermate control). A total of 31 Cre positive (+) mice were collected at 8 weeks of age, 7 male IFT88^+/+^ (WT), 8 male IFT88^LoxP/LoxP^ (cKO), 8 female IFT88^+/+^ (WT), and 8 female IFT88^LoxP/LoxP^ (cKO).

Dmp1-8kb-Cre mice possessing the transgene orienting the Cre recombinase sequence driven by the murine dentin matrix protein 1 (Dmp1) gene [22] were bred with MKS5^LoxP^ mice, generating Dmp1-Cre; MKS5 ^+/LoxP^ mice that were crossed with the purpose of producing Dmp1Cre; MKS5^LoxP/LoxP^ (cKO) mice and WT controls, Dmp1-Cre; MKS5^+/+^ [21, 23], in order to obtain mice with osteocyte-specific deletion of the MKS5 gene. A total of 29 Cre (+) mice were collected at 20 weeks of age, 8 male MKS5^+/+^ (WT), 8 male MKS5^LoxP/LoxP^ (cKO), 7 female MKS5^+/+^ (WT), and 6 female MKS5^LoxP/LoxP^ (cKO).

The genotype of each mouse was determined and confirmed by polymerase chain reaction (PCR) amplification of tail DNA samples at weaning and sacrifice. All primers were acquired through IDT (Integrated DNA Technologies, Inc., Skokie, IL, USA). When genotyping for Osx-Cre, the primer sequences used for its amplification were as follows: Osx-cre forward (TGCK) – 5’ - CTC TTC ATG AGG AGG ACC CT -3’, Osx-cre reverse (OSX) – 5’ - GCC AGG CAG GTG CCT GGA CAT -3’. For the amplification of IFT88 LoxP, the primers used were IFT88 Common forward – 5’ – GCC TCC TGT TTC TTG ACA ACA GTG - 3’, IFT88 LoxP and WT Reverse – 5’ – GGT CCT AAC AAG TAA GCC CAG TGT T - 3’, and IFT88 Null Reverse – 5’- CTG CAC CAG CCA TTT CCT CTA AGT CAT GTA - 3’. When genotyping for Osx-Cre, a 615-bp band indicated the presence of the recombinase, while the lack of a band indicated its absence. When genotyping for IFT88 LoxP, a 350-bp band indicated IFT88^flox/flox^ individuals, while a 310-bp band signified IFT88^+/+^ individuals. Heterozygous individuals were signified by the presence of three bands, at 350-bp, 310-bp, and 280-bp.

The primer sequences used for the amplification of Dmp1-Cre were Dmp1-Cre forward (CsCre1) – 5’ - AGG GAT CGC GAG GCG TTT TC - 3’ and Dmp1-Cre reverse (CsCre2) – 5’ - GTT TTC TTT TCG GAT CCG CC -3’. For the amplification of MKS5 LoxP, there were three primer sequences used. They were MKS5 Common forward – 5’ - GTC CTC TGA CTT CCA GTG TCA TGT GC - 3’, MKS5 LoxP and WT Reverse – 5’ - GTG GGT TGT ACA GTT TCT GCT TCA TCC AC - 3’, and MKS5 Null Reverse – 5’ - AAG CTC TAA AGC TGG GAC TGC AGC - 3’. When genotyping for Dmp1Cre, the presence of a 300-bp band indicated the presence of Dmp1-Cre recombinase, while the lack of a band indicated its absence. When genotyping for MKS5 LoxP, one single 788-bp band indicated MKS5^flox/flox^, while one single 605-bp band indicated MKS5^+/+^. The presence of both bands together indicated heterozygous individuals with only one LoxP site. The heterozygous mice from both the MKS5 and the IFT88 lineages were only used for breeding purposes.

### 3.2 In Vivo Ulnar Loading

In vivo axial ulnar loading was performed on 20-week-old (± 14 days) Dmp1-Cre; MKS5^LoxP^ mice and their littermate controls. The animals were subjected to axial ulnar loading for 3 consecutive days as previously published [24]. The right ulnae were loaded for 120 cycle/day at a frequency of 2Hz with a peak force of 2.9 N for female mice and 3.2 N for male mice. The left ulnae served as internal controls. All animals were allowed regular cage activities following loading sessions. Two days following the final day of loading (10 days prior to sacrifice), the animals were injected with 0.4 mL green fluorochrome label calcein (30 mg/Kg body weight) via intraperitoneal injection, and 7 days later (3 days prior to sacrifice) they were administered 0.4 mL of red fluorochrome label alizarin (50 mg/Kg body weight) via intraperitoneal injection. All test animals were sacrificed 3 days after the administration of alizarin.

### 3.3 Histology and Histomorphometry

For the Osx-Cre; IFT88 test groups, the right femur was processed for histological and histomorphometric analysis. For the Dmp1-Cre; MKS5 test groups, the right distal femur and right and left ulnae were processed for histological and histomorphometric analysis. All samples were processed following the protocol previously established [24].

The bones were placed in a vial containing 10 mL of 10% neutral-buffered formalin (NBF) and stored in 4°C for 24 hours. The formalin was then replaced by 70% EtOH and placed in the 4°C storage until ready to be processed. The bones were dehydrated by a series of graded alcohols with the concentration increased every 2-8 hours until it reached 100% EtOH. The 100% EtOH was replaced by a 1:1 mixture of EtOH and methyl methacrylate (MMA) for 2-8 hours. Then the samples were placed in undiluted MMA for 2-8 hours, which was later replaced by infiltration media (MMA and 4% dibutyl phthalate) and placed under vacuum for 2-7 days at 17 inHg.

Using a wire saw, the ulna was cut proximal from the pencil mark drawn at the midshaft during tissue processing to ensure midshaft collection. Then 5 sequential transverse thick sections (50μm) were collected. The sections were mounted on microscope slides using Eukitt^®^ mounting medium (MilliporeSigma, St. Louis, MO, USA) and xylenes-thinned mounting medium were applied to the mounted section, increasing the surface area for grinding. The slides were ground to 30-50 μM against wet 600 grit sandpaper, then they were cover slipped using xylenes-thinned mounting medium.

MMA blocks embedding the distal femur were cut into thin sections on a rotary microtome (Leica Biosystems Inc., Buffalo Grove, IL) using a tungsten carbide knife. The block was secured on the microtome and 50% EtOH was applied to the face of the block for softening. Five microscope slides previously subbed with 1% gelatin solution (Thermo Fisher Scientific, Waltham, MA, USA) were prepared, each containing three 4 μm thick sections. The slides were left to dry at 37°C overnight, then cover slipped.

Quantitative analysis of bone samples was performed using OsteoMeasure™ histomorphometry software (OsteoMetrics, Decatur, GA, USA). The slides were analyzed using an X-Cite Series 120Q fluorescence illuminator (Excelitas Technologies, Waltham, MA, USA) on an Olympus DP72 digital microscope camera connected to an Olympus BX53 Telepathology Microscope System (Olympus Scientific Solutions Americas, Waltham, MA, USA). Measurements were taken by tracing the sections at 200x magnification on a touchscreen (Wacom, Kazo, Saitama Prefecture, Japan).

Unstained cortical sections of the left and right ulnae, as well as an unstained right femur midshaft cortical section of each study subject were analyzed for periosteal perimeter, endocortical perimeter, periosteal single label, endocortical single label, periosteal double label, and endocortical double label. Using these tracings, derived measurements, such as mineralizing surface (MS/BS), mineral apposition rate (MAR), and bone formation rate (BFR/BS), were calculated. As the left ulna was used as a non-loaded control, the values for the left ulna were subtracted from the values for the right ulna, resulting in the amount of bone activity occurring because of loading.

One slide of the distal femur of each test subject was stained with tartrate-resistant acid phosphatase (TRAP) and counterstained with a toluidine blue. Histomorphometry was performed using OsteoMeasure™ histomorphometry software (OsteoMetrics, Decatur, GA, USA). The slides were analyzed under bright-field illumination on an Olympus DP72 digital microscope camera connected to an Olympus BX53 Telepathology Microscope System (Olympus Scientific Solutions Americas, Waltham, MA, USA). Each sample was analyzed for osteoclast number (N.Oc), osteoclast surface (Oc.S/BS), osteoclast number over bone surface (N.Oc/B.Pm), and osteoclast number over perimeter (N.Oc.Oc.Pm).

### 3.4 Peripheral Dual-Energy X-Ray Absorptionmetry (PIXImus)

The bone mineral density (BMD, g/cm^2^) and mineral content (BMC, g) of the left femur of each test animal were analyzed using a PIXImus dual-energy X-ray absorptiometer (DXA) (PIXImus Lunar Corp., Madison, WI, USA).

### 3.5 Micro-Computed Tomography (μCT)

The left femur of each test animal was imaged with a SkyScan 1172 microCT (SkyScan, Kontich, Belgium) following the protocol previously established [25]. Reconstruction of the 3D structures was done through NRecon reconstruction software (Micro Photonics Inc., Allentown, PA, USA). CTan software was used to select ROI for each midshaft. Cortical ROIs composed of 5 images each, averaging 10 μm of cortical bone surface was utilized in this analysis. The trabecular ROI was defined as 1 mm of the trabecular region of the distal femur beginning 1 mm proximal to the growth plate.

### 3.6 Biomechanical Testing

The left femora were tested in three-point bending tests to failure to evaluate structural-level and tissue-level mechanical properties. The support span was set at 5.4mm and 8 mm for Osx-Cre;Ift88 and Dmp1-Cre;MKS5, respectively. Each femur was oriented so the loading point was applied at the 50% region of the bone and the anterior surface of the bone was in tension. Each test was conducted under displacement control at a loading rate of 0.025 mm/sec until failure. During each test, the resulting force and displacement were recorded.

Structural-level mechanical properties were obtained from collected force-displacement data and included ultimate force, yield force, total work and stiffness. To calculate tissue-level mechanical properties of the bones, the recorded force and displacement data were converted to stress and strain using geometric properties obtained from µCT analysis. A region of interest of 10 transverse slices was obtained from µCT scans at the 50% region of each bone and used to calculate the necessary geometric properties. These geometric properties were then used along with the force and displacement data in a custom MATLAB (MathWorks, Natick, MA) script, from Professor Joseph Wallace at Indiana University-Purdue University Indianapolis, Department of Biomedical Engineering (BME), to calculate stress and strain using standard engineering equations to then obtain tissue-level mechanical properties. The collected structural-level mechanical properties included ultimate stress, Young’s modulus and toughness [26].

### 3.7 Imaging of Primary Cilia

Serial digestion of calvariae from newborn Osx-Cre;Ift88 mice (1-5 days old) was performed to isolate primary bone cells. Calvariae were dissected and cleaned from any soft tissue and placed in media (α-MEM, 10% FBS, 1% P/S) while PCR and gel electrophoresis were run to confirm the calvaria were of the correct genotype. Once the genotype had been determined, the calvaria were cut into 3-4 pieces and placed in conical tubes with 3 mL of digest solution, composed of 0.1% collagenase, 0.05% trypsin, 4 mM EDTA. The tubes were then placed on a shaker for 20 minutes. The supernatant was removed, filtered, and transferred to another conical tube. This process was repeated 4 times, and after the last one, the new conical tubes containing the supernatant were centrifuged. The cells were plated and left to grow.

Once the cells were confluent, they were starved in media containing α-MEM, 0.1% FBS and 1% Penicillin-Streptomycin antibiotic (P/S) for 24 hours to promote primary cilium growth. Cells were fixed with 4% PFA 10% sucrose solution for 10 minutes at room temperature. The solution was then aspirated, the cells washed with 1x PBS, and 100% Methanol was added to the cells for 20 minutes. Once the Methanol was aspirated off, 1x PBS with 0.1% Triton X was added and the plate was incubated for 6 minutes. The cells were then washed with 1x PBS again, and a blocking solution (PBS+, composed of 2% donkey serum, 0.3% Triton X-100, 1% Bovine Serum Albumin, 10X PBS, and 0.02% Sodium Azide) was added after. The cells were left in room temperature with the blocking solution for one hour. The primary antibody (Monoclonal Anti-Acetylated Tubulin, Sigma T7451) was then added, and the cells were incubated overnight. The cells were once again washed with PBS+, and the secondary antibody (Alexa Fluor™ 488 donkey anti-mouse IgG (H+L), Invitrogen #A21202) was added following the wash for one hour. The cells were then washed in PBS+ and Hoechst (Hoechst 33342, ThermoFisher Scientific #62249) was added in a 1:1000 dilution. The cells were then ready to be imaged.

### 3.8 Statistical Analysis

The data are expressed as mean ± SD (standard deviation). The data were analyzed and compared using One-Way ANOVA using GraphPad Prism (GraphPad Software, San Diego, CA). Significance was determined as p<0.05.

## 4. Results

### 4.1 Primary cilia in osteoblasts affect bone growth in mice

#### 4.1.1 Body Weight, Bone Size, and Density

A significant decrease in body weight was observed in both male (p=0.0048) and female (p=0.0374) groups when comparing conditional knockout mice to wild type controls. The average body weights in grams (g) with their standard deviations are as follows: WT male 20.51 ± 1.168, cKO male 18.42 ± 1.1909, WT female 17.736 ± 2.796, cKO female 15.655 ± 1.824 (Figure 1A & B).

**Figure 1.**
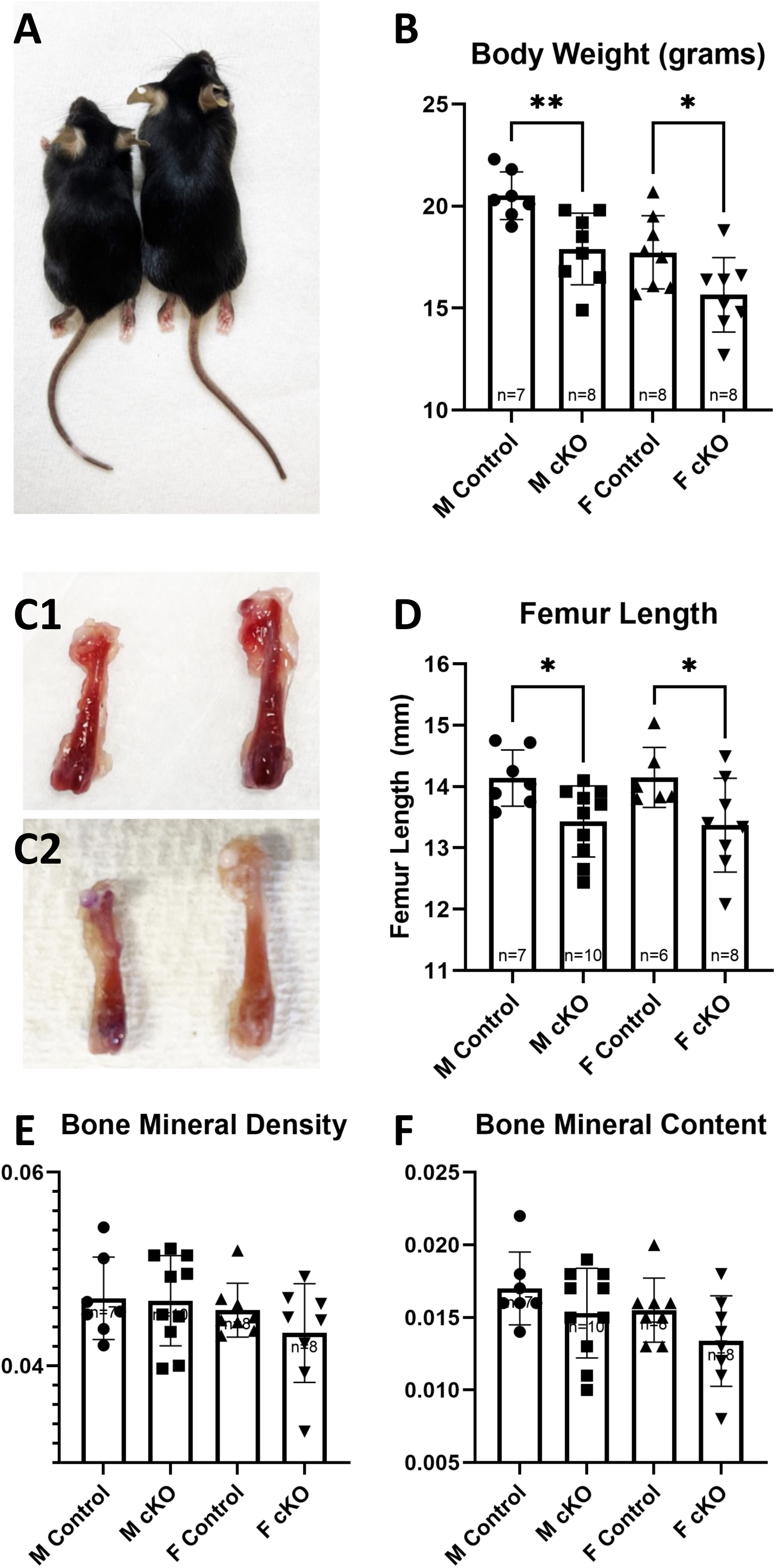
Body weight, femoral length, BMD and BMC of Osx-Cre; Ift88 mice. (A) Photo example of difference in size between WT and cKO mice. (B) Body weights of WT and cKO mice at 8 weeks of age. Animals of cKO groups had significantly lower body weights compared to WT animals. (C1) Photo example of difference in length of femur between male cKo (Left) and WT (Right). (C2) Photo example of difference in length of femur between female cKo (Left) and WT (Right). (D) Length of femur of WT and cKO mice at 8 weeks of age. Male and female cKO had significantly shorter femurs when compared to WT. Values for (E) BMD and (F) BMC. PIXImus scans of femurs reveal no difference in either BMD or BMC of cKO animals compared to WT. Note that *: p < 0.05, **: p<0.01.

The femurs of conditional knockout mice were significantly shorter than the femurs of wild type mice of both male (p=0.0003) and female (p=0.0019) groups. The average length of femurs in millimeters (mm) with their standard deviations are as follows: WT male 1.457 ± 0.053, cKO male 1.31 ± 0.074, WT female 1.428 ± 0.0488, cKO female 1.3 ± 0.0756 (Figure 1 C1, C2 & D). The bone mineral content (BMC) and bone mineral density (BMD) of cKO mice was slightly lower than that of the WT mice, but no significance was reached (Figure 1E & F). This data suggests primary cilia in osteoblasts affect bone growth in mice.

Immunofluorescent staining confirmed that cilia was knocked out from osteoblast cells (Figure 2). In the WT cells, primary cilia can be seen, whereas they are gone from the cells of cKO mice.

**Figure 2.**
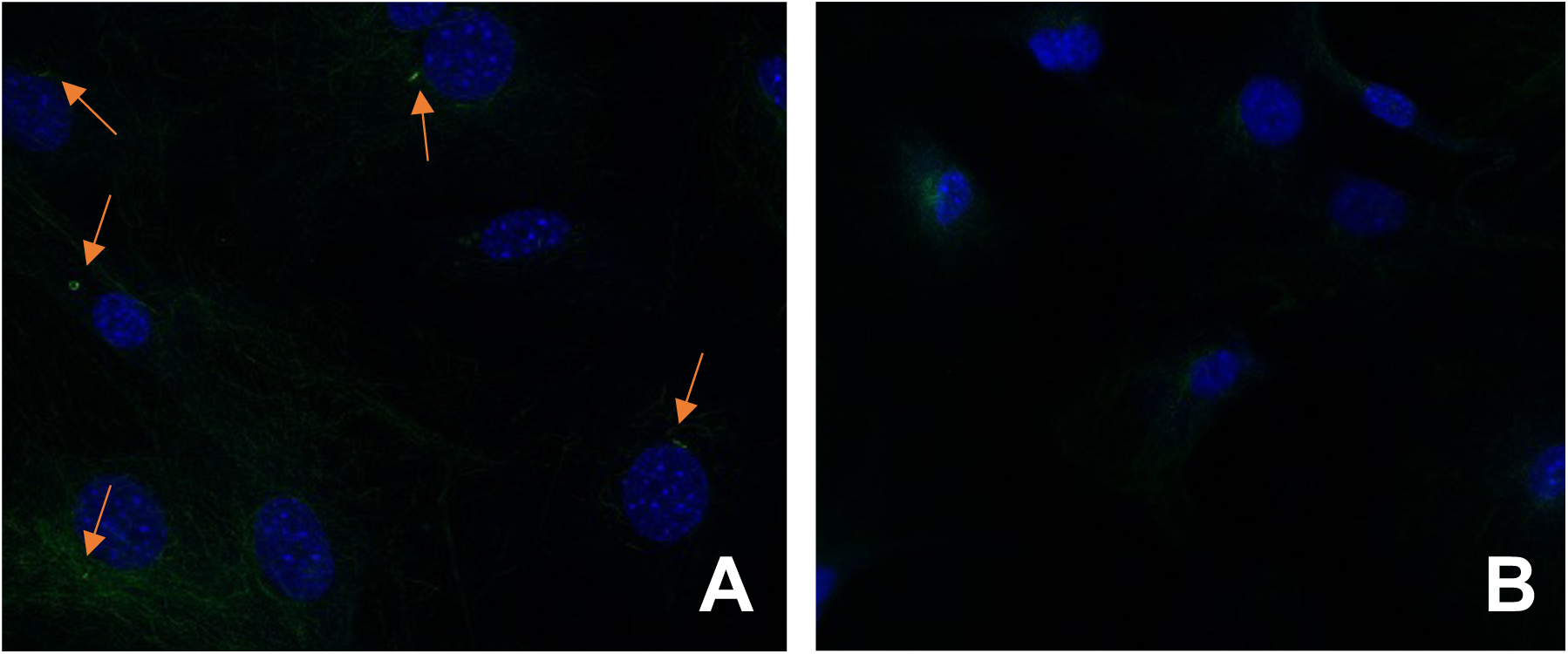
Representative images of cilia in (A) in OsxCre; IFT88^+/+^ and (B) OsxCre; IFT88^LoxP/LoxP^.

#### 4.1.2 Trabecular Bone: Structure, Bone Formation and Resorption

Trabecular analysis of Micro-CT scans found that female cKO mice had a significant increase in trabecular number (p=0.0443) when compared to female WT mice. Other measured parameters, including bone volume, BV/TV, bone surface, and thickness showed a non-significant trend in which both male and female cKO had favored an increase compared to WT controls (Figure 3).

**Figure 3.**
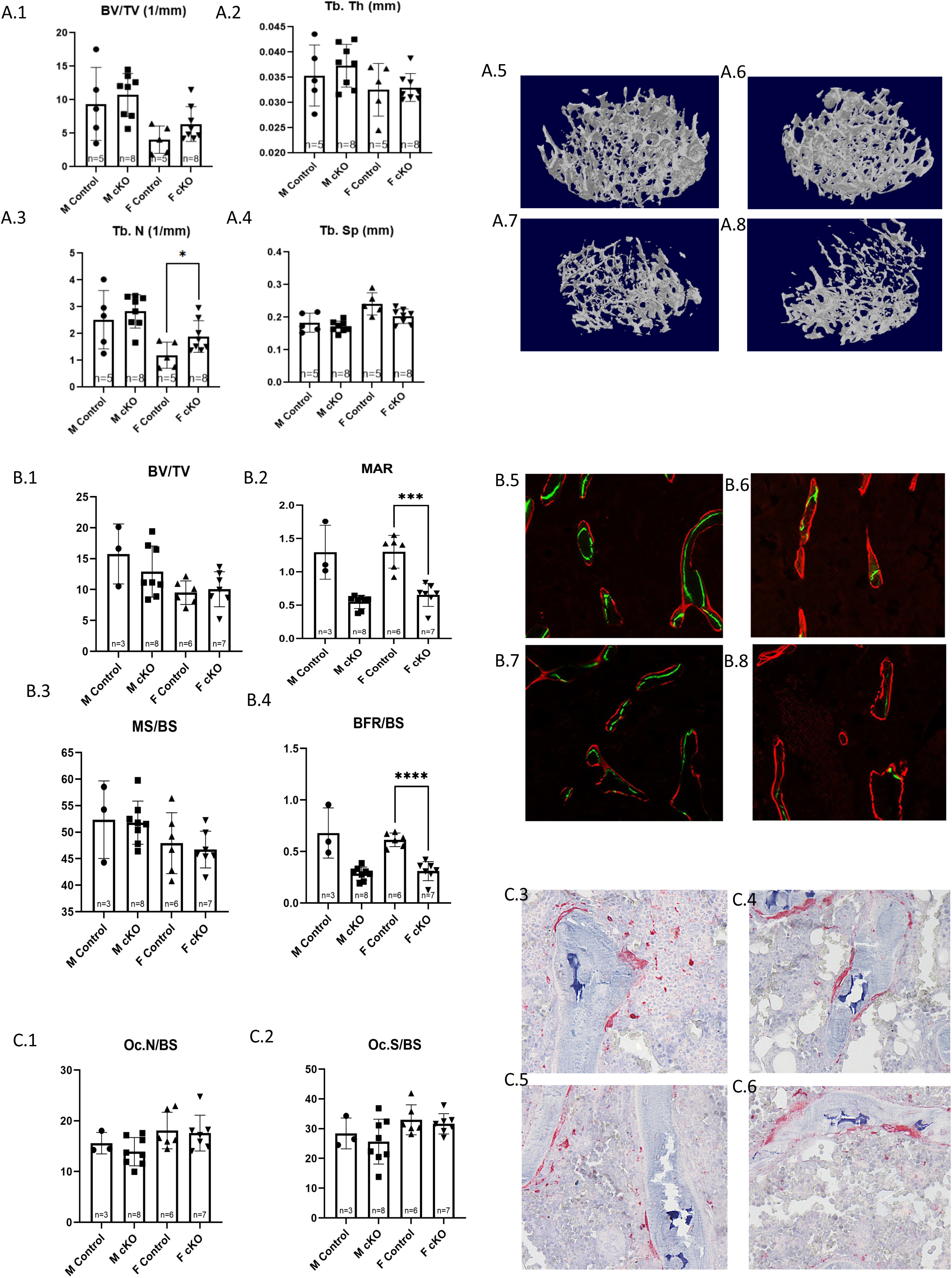
Trabecular analysis of Osx1Cre^+^, IFT88^LoxP/LoxP^ (cKO) Mice. (A1-4) Trabecular analysis results for distal femur of male and female cKO mice and WT controls; accompanied by 3D reconstructions of trabeculae from (A5) male WT, (A7) male cKO, (A6) female WT, and (A8) female cKO mice. **Dynamic Histomorphometry Analysis of Distal Femur Trabeculae in OsxCre; IFT88^LoxP/LoxP^** was done for (B1) Trabecular BV/TV, (B2) Trabecular MAR, (B3) Trabecular MS/BS, and (B4) Trabecular BFR/BS. Measurements of trabecular MAR and BFR of the female cKO mice was significantly lower than that of the WT females. Photo examples show trabecular fluorescent labeling in (B5) WT male, (B7) cKO male, (B6) WT female, and (B8) cKO female mice. TRAP-stained thin trabecular sections of the distal femur of OsxCre; IFT88^LoxP/LoxP^ (cKO) mice were compared to those of wild-type (WT) controls. No significant difference was observed in any of the measured parameters, including (C1) Osteoclast number proportional to bone surface and (C2) Osteoclast surface proportional to bone surface. Photo examples show TRAP stain in (C3) WT male, (C5) cKO male, (C4) WT female, and (C6) cKO female mice. Note that *: p < 0.05, ***: p < 0.001, ****: p<0.0001.

Histological analysis of the thin trabecular sections revealed a significant difference in MAR (p=0.0005) and BFR/BS (p<0.0001) between female WT and cKO mice. Although it did not reach significance, the male conditional knockout group had a MAR that was 57.57% lower and a BFR/BS rate that was 58.03% lower compared to the wild-type control group (Figure 3).

TRAP positive osteoclasts were counted on thin sections of the distal femur to estimate bone resorption. When compared between WT and cKO groups, no significance in osteoclast surface proportional to bone surface or osteoclast number proportional to bone surface was observed (Figure 3).

Mechanical testing of the femur in three-point bending test showed no differences in structural mechanical properties between femurs of cKO and WT groups (Figure 4). Whereas there were no significant differences in the ultimate force and stiffness, the average of the cKO groups were generally lower than that of the WT groups. Young’s modulus was significantly higher in the male cKO group compared to the WT group.

**Figure 4.**
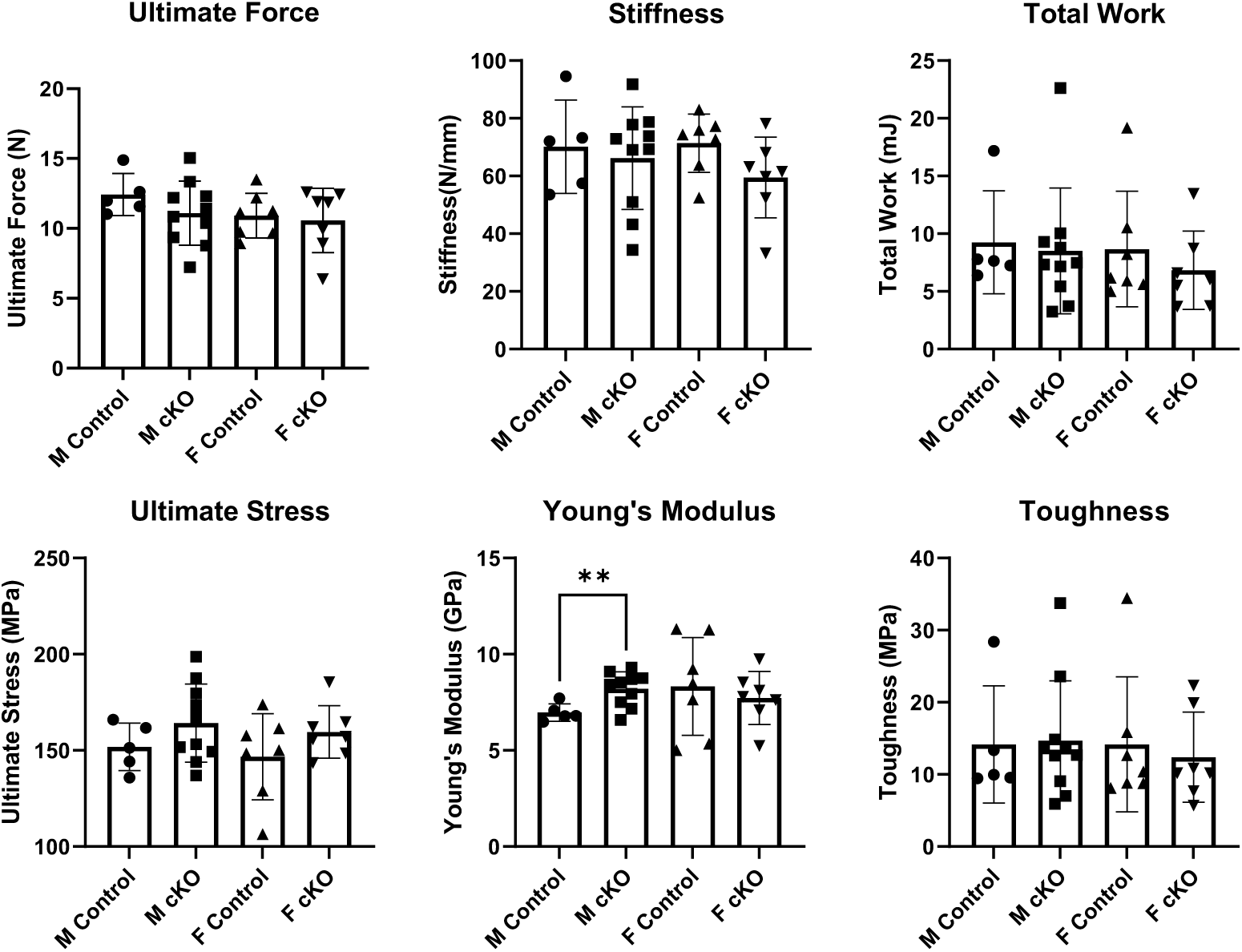
Three-Point bending analysis of Osx-Cre; Ift88 mice.

**Figure 5.**
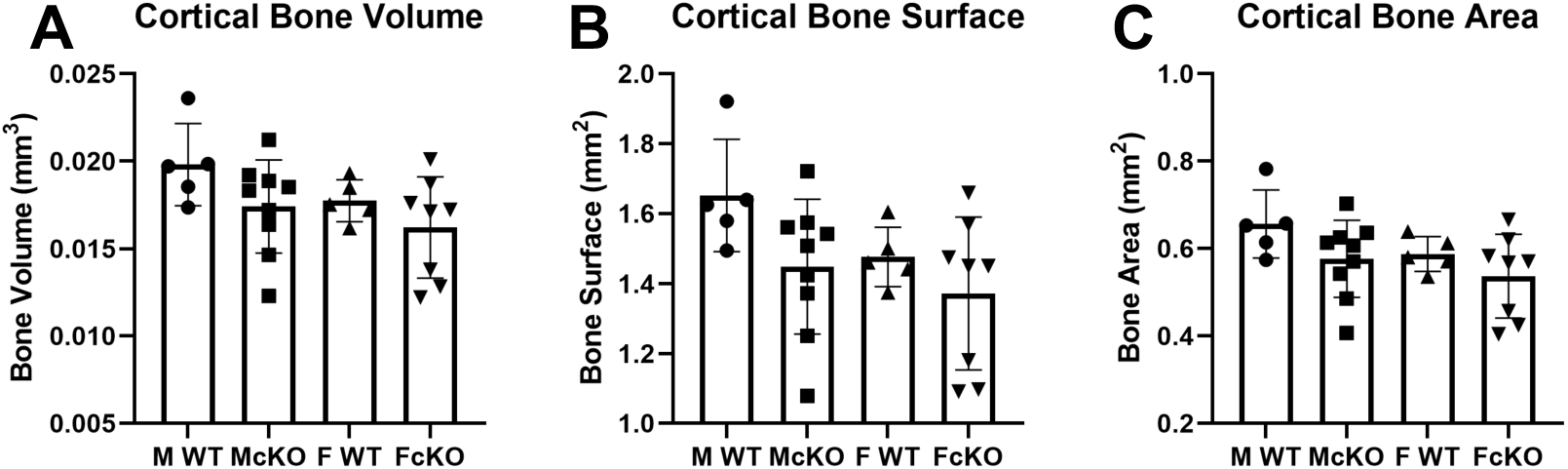
Cortical analysis of OsxCre^+^, IFT88^LoxP/LoxP^ (cKO) Mice. Results collected from left femora of male and female cKO and WT controls.

#### 4.1.3 Cortical Bone: Structure and Bone Formation

Cortical analysis of femoral midshafts found no significant difference between WT controls and cKO mice in bone volume, bone surface, or bone area. It was observed, however, that both the male and female cKO groups showed a non-significant trend towards a decreased bone area when compared to their WT controls (Figure 9).

### 4.2 Primary cilia in osteocytes play an important role in mechanotransduction

#### 4.2.1 Body Weight, Bone Size and Density

A significant decrease in body weight between male WT and male cKO mice was observed (p=0.0011). Although the same trend was observed on the female mice, it did not reach significance. The average body weights with their standard deviations are as follows: WT male 28.75 ± 1.13641, cKO male 26.3625 ± 1.197542, WT female 24.64286 ± 2.053336, cKO female 23.38333 ± 1.725592 (Figure 6A). Male cKO mice exhibited a trend toward shorter femurs than control mice but did not reach the level of statistical significance (p=0.0537). No difference was observed within the female groups (Figure 6B, C1 & C2) Bone mineral density and mineral content were analyzed using PIXImus. Although a trend was observed in which the bone mineral content (BMC) and bone mineral density (BMD) of cKO mice was slightly lower than that of the WT mice, no statistical significance was reached in either male or female groups (**Error! Reference source not found.** 6D & E).

**Figure 6.**
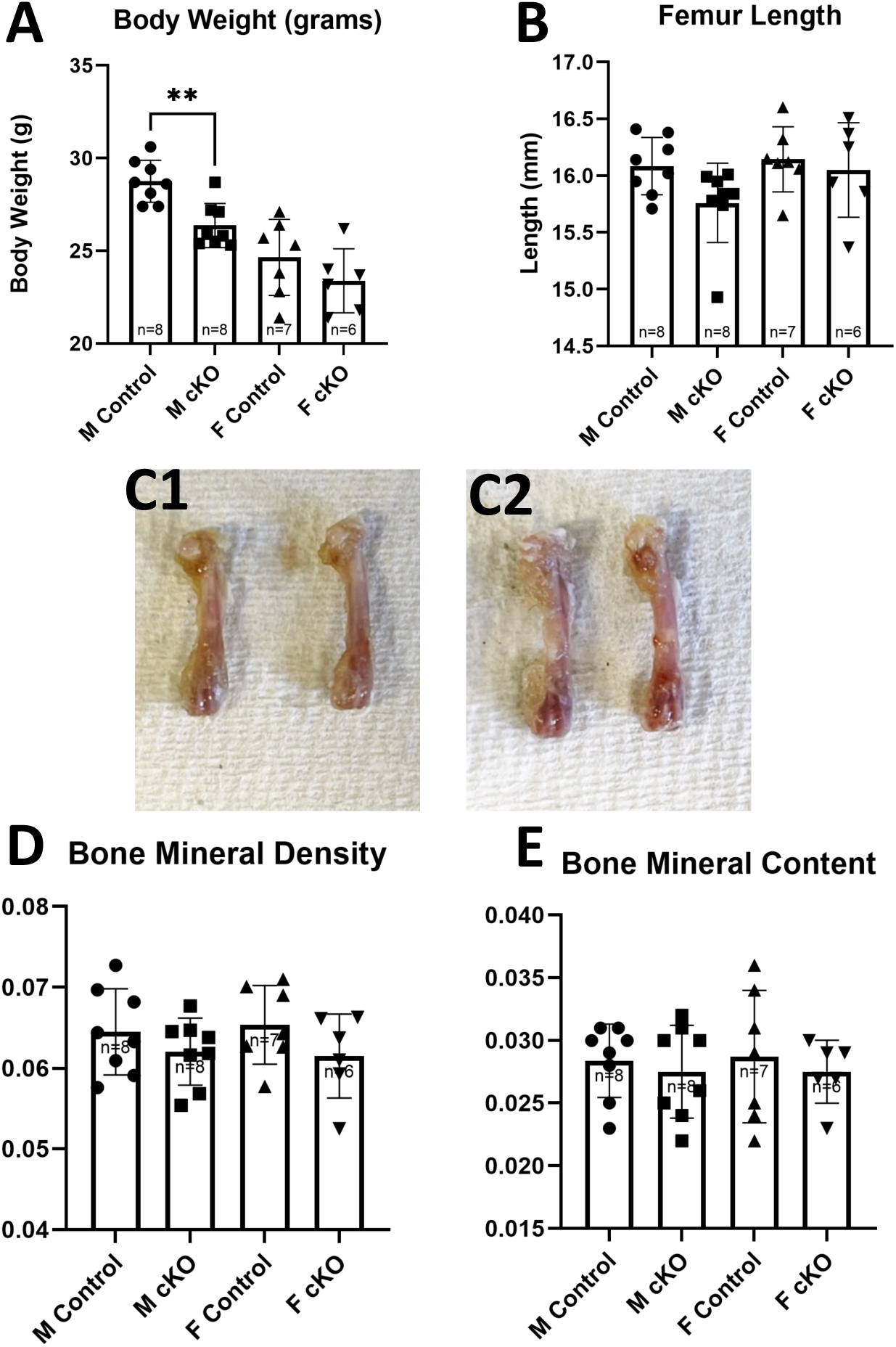
Body weight, Length of femur, BMD and BMC of Dmp1Cre; MKS5 mice. (A) Body weights and (B) length of femur of WT and cKO mice at 20 weeks of age. Male cKO mice had significantly lower body weights compared to WT males. (C1) Photo of femur of male cKO (Left) and WT (Right) mice. (C2) Photo of femur of female cKO (Left) and WT (Right) mice. Values for (D) BMD and (E) BMC. PIXImus scans of femurs reveal no difference in either BMD or BMC of cKO animals compared to WT. Note that **: p < 0.01.

#### 4.2.2 Trabecular Bone: Structure, Formation, and Resorption

Trabecular analysis of micro-CT scans showed no significant differences in BV/TV in either male or female groups. It found that female cKO mice had a significant decrease in trabecular number (p=0.0207) and trabecular bone surface (p=0.0497) when compared to female WT mice, as well as a significant increase in trabecular separation (p=0.0313). Male cKO mice exhibited significantly decreased trabecular thickness (p=0.0419) compared to their WT controls (Figure 7).

**Figure 7.**
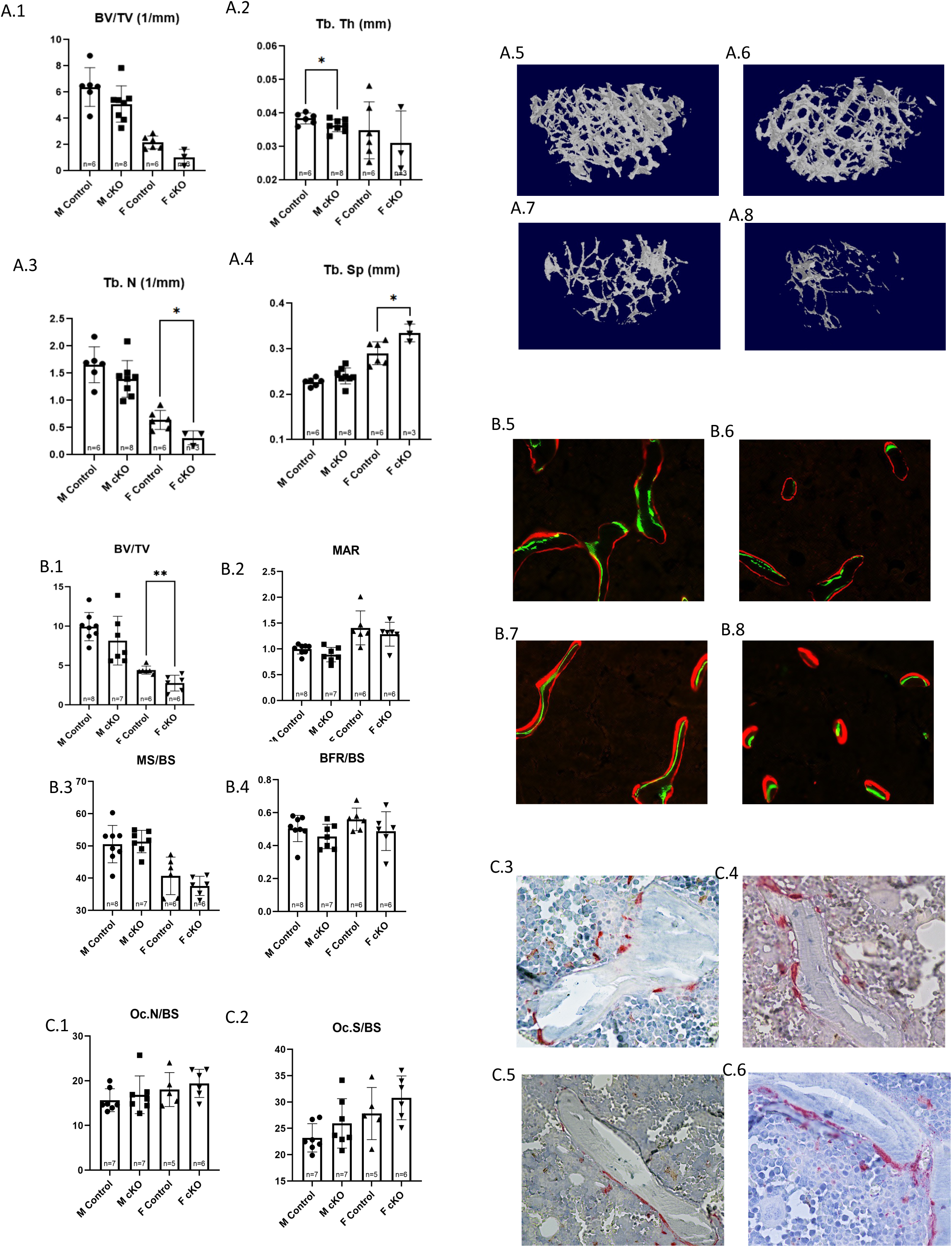
Trabecular analysis of Dmp1Cre^+^, MKS5^LoxP/LoxP^ (cKO) Mice. (A1-4) Trabecular analysis results for distal femur of male and female cKO mice and WT controls; accompanied by 3D reconstructions of trabeculae from (A5) male WT, (A7) male cKO, (A6) female WT, and (A8) female cKO mice. **Dynamic Histomorphometry Analysis of Distal Femur Trabeculae in Dmp1-Cre; MKS5^LoxP/LoxP^** was done for (B1) Trabecular BV/TV, (B2) Trabecular MAR, (B3) Trabecular MS/BS, and (B4) Trabecular BFR/BS. Measurements of trabecular BV/TV of the female cKO mice was significantly lower than that of the WT females (p=0.008). Photo examples show trabecular fluorescent labeling in (B5) WT male, (B7) cKO male, (B6) WT female, and (B8) cKO female mice. TRAP-stained thin trabecular sections of the distal femur of Dmp1-Cre; MKS5^LoxP/LoxP^ (cKO) mice were compared to those of wild-type (WT) controls. No significant difference was observed in any of the measured parameters, including (C1) Osteoclast number proportional to bone surface and (C2) Osteoclast surface proportional to bone surface. Photo examples show TRAP stain in (C3) WT male, (C5) cKO male, (C6) WT female, and (C8) cKO female mice. Note that *: p < 0.05, **: p < 0.01.

Histological analysis of the thin trabecular sections revealed a significant difference in BV/TV between WT and cKO females (p=0.008). Although it did not reach significance, male conditional knockout mice had a BV/TV that was 17.93% lower than that of wild type males. Even without reaching significance, a trend was noticed in all other parameters (MAR, MS/BS, and BFR/BS) in which the cKO values were lower than that of WT mice for both sexes (Figure 7). These data suggest that primary cilia in osteocytes influence bone formation in trabecular bone.

TRAP staining positive osteoclasts were counted on thin sections of the distal femur of all the animals. No significances in osteoclast number and osteoclast surface were observed when compared between WT and cKO groups. Osteoclast surface proportional to bone surface and osteoclast number proportional to bone surface were the measured parameters (Figure 7). The male knockout mice had an increase of 11.85% on osteoclast surface compared to wild type males, while female conditional knockout mice had an increase of 4% compared to the wild type females. The number of osteoclasts present on stained thin sections of the male knockout mouse was 7.73% higher than that of the male wild-type mouse, and for the female knockout mouse, it was 1.05% higher than the female wild type. These data suggest that primary cilia in osteocytes do not influence the effects of osteocytes on osteoclasts.

Analysis of the three-point bending test showed no significant differences in most parameters measured (Figure 8). Total work required was significantly less for male cKO group compared to their sex-matched control, and although the average of the female cKO group showed a lower trend compared to the female control group, no significance was reached. Both male and female cKO groups also showed a trend towards lower toughness when compared to their respective controls, although no significance was reached in that parameter either.

**Figure 8.**
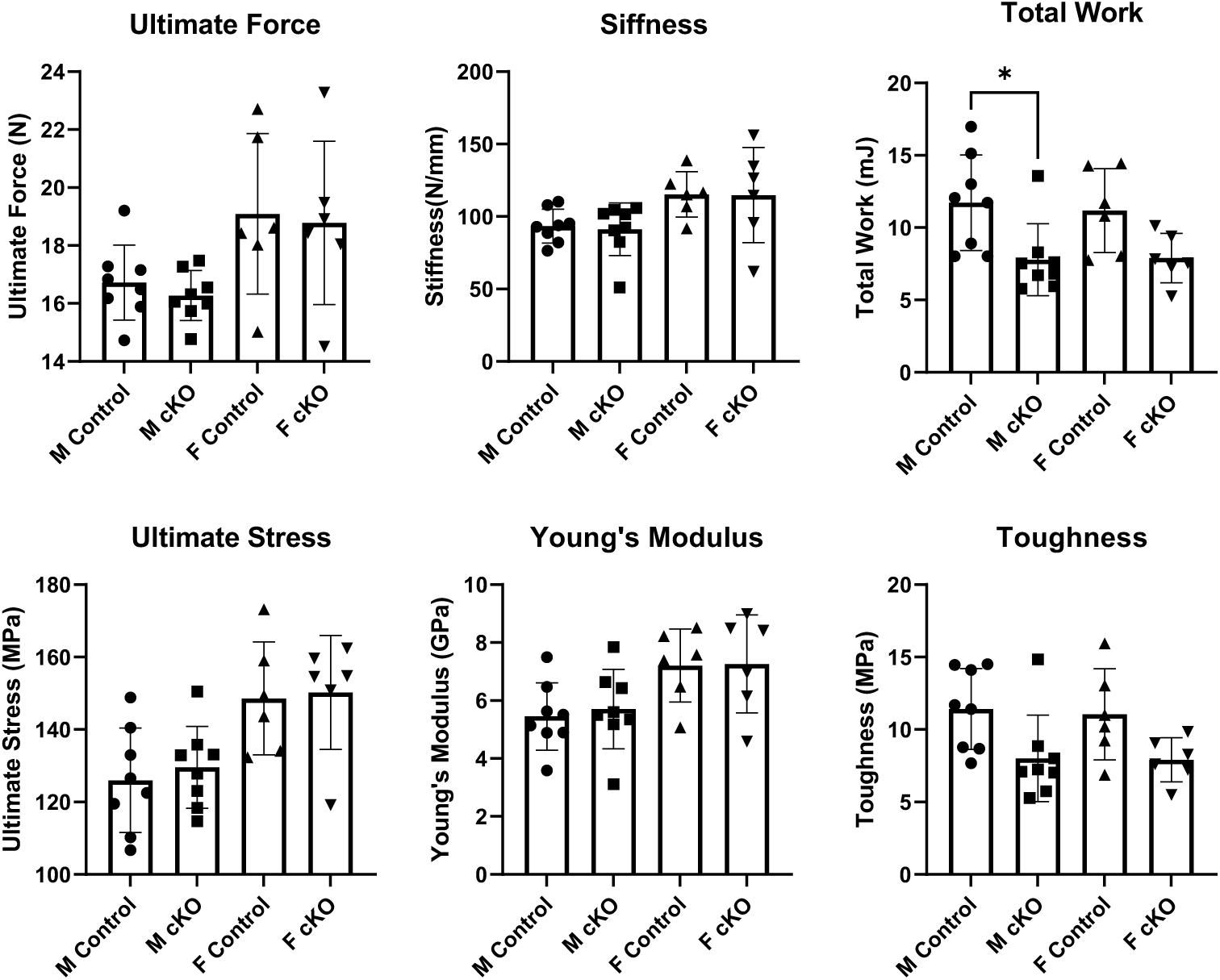
Three-Point bending analysis of Dmp1-Cre; MKS5 mice.

#### 4.2.3 Cortical Bone: Structure and Bone Formation

Cortical analysis of micro-CT scans found no significant difference between WT controls and cKO mice in any of the measured parameters, including bone volume, bone surface, bone marrow area, and bone area (Figure 9).

**Figure 9.**
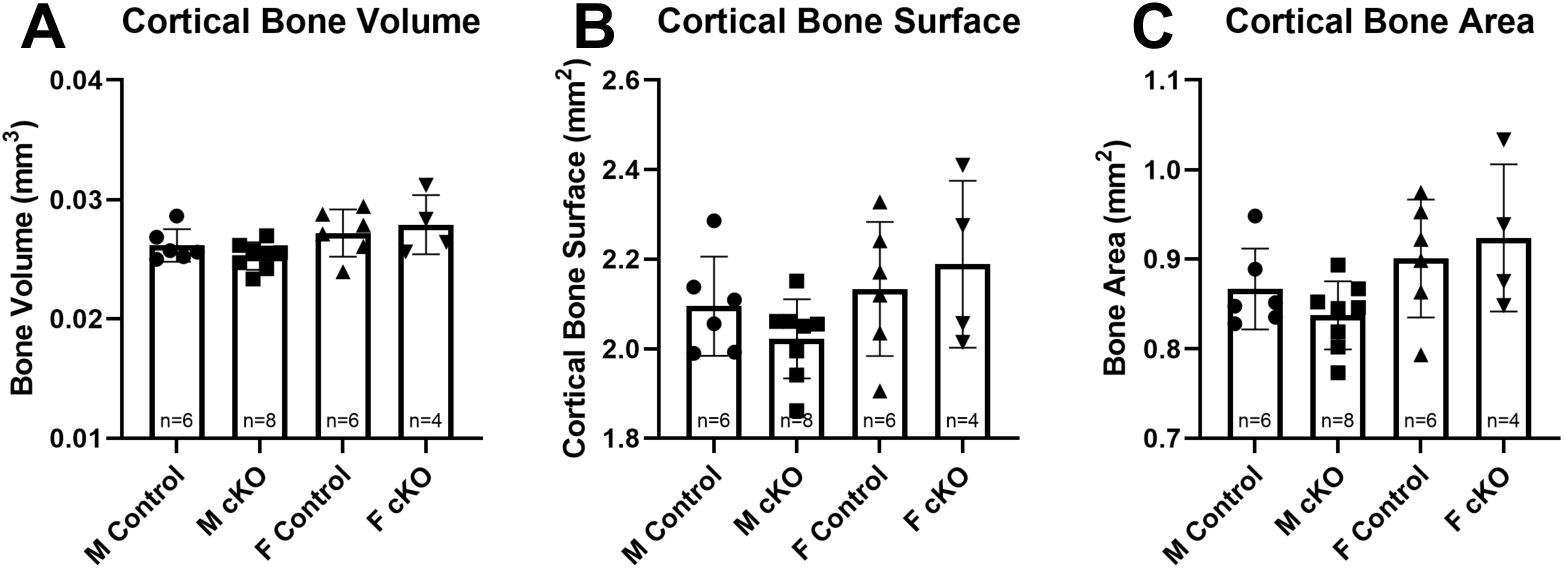
Cortical analysis of Dmp1Cre^+^, MKS5^LoxP/LoxP^ (cKO) Mice. Results collected from left femora of male and female cKO and WT controls.

**Figure 10.**
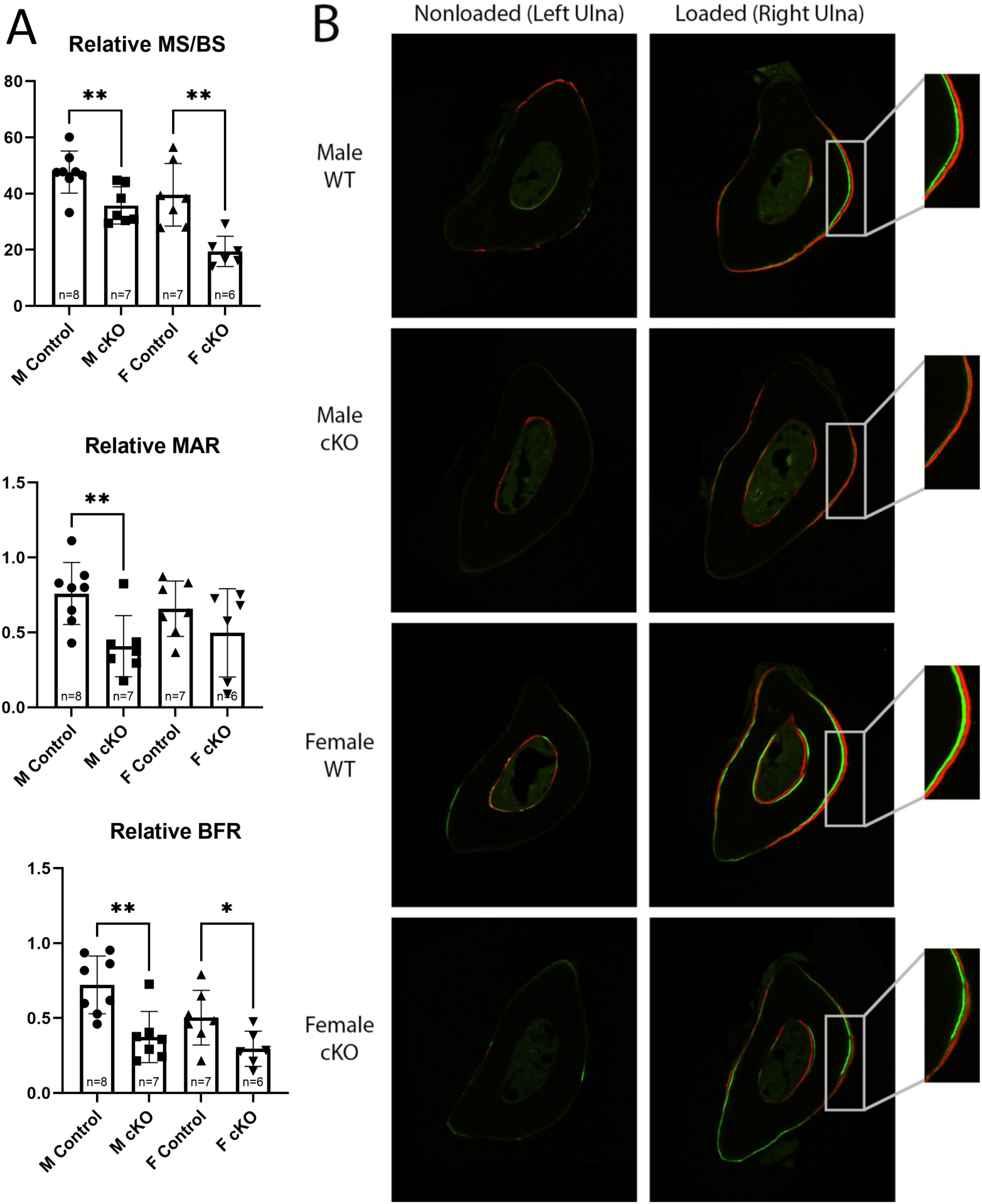
Loading-Induced Bone Formation of Dmp1Cre^+^, MKS5^LoxP/LoxP^ (cKO) Mice. (A) The relative values, calculated as the difference between the right (loaded) and left (non-loaded) ulna was calculated in order to determine bone formation activity in the ulna as a result of mechanical loading. (B) Photo examples of cortical sections of right and left ulna from all four test groups. Note that *: p < 0.05, **: p<0.01.

#### 4.2.4 Mechanically Induced Bone Formation

Histological analysis of the cortical sections of the left and right ulna was performed to determine loading-induced bone formation. Significant differences in relative MS/BS and relative BFR/BS between male and female WT and cKO were observed. The relative MAR parameter only revealed a significant difference between male WT and cKO mice (p=0.0058), although a slight, though non-significant, difference was observed between the female WT and cKO groups. Within the male groups, the conditional knockout mice had a MS/BS (p=0.0064) that was 24.88% lower than that of the control group, as well as a MAR (0.0058) that was 46.27% lower and BFR/BS (p=0.0026) that was 48.24% lower. For the female groups, the MS/BS (p=0.0022) was 52.5% lower for the knockout group compared to the control. The MAR was 27.58% lower, and the BFR/BS (p=0.0325) was 41.54% lower when compared to the wild type control group (Figure 18). In summary, this data suggests primary cilia in osteocytes play an important role in load induced bone formation.

## 5. Discussion

This study examined the mechanosensory role of primary cilia in bone growth and adaptation by examining two cilia specific genes, IFT88 and MKS5, required for proper cilia assembly and function. The results of this study indicate that the osteocyte primary cilium affects bone homeostasis and mechanotransduction, presenting evidence that primary cilia act as mechanosensors in bone cells.

Conditional knockout (cKO) mice appeared smaller than their wild-type (WT) controls. This observed phenotype was significant in males, and although the same patterns could be observed in females, significance was not reached. Male cKO body weights at 20 weeks of age were significantly lower than WT controls, showing a trend toward shortened femur lengths as well. Despite lower body weights and a seemingly smaller femur, no significant differences were observed in BMD and BMC of either sex when compared to their WT counterparts. In trabecular sections, the female cKO group exhibited a significant decrease in trabecular BV/TV compared to WT controls, and although their MS/BS, MAR, and BFR were lower as a group, it never reached significance. The cKO male group had an overall lower BV/TV, MAR, and BFR/BS than the male WT group; however, none of the differences observed were significant. These differences, though non-significant in this study, could indicate slightly decreased osteoblastic activity in cKO mice compared to that of WT mice. No significant differences were observed in osteoclast number or osteoclast surface. Dynamic ulnar loading showed a striking change in mechanotransduction when comparing cKO to WT. Both male and female cKO mice had significantly lower relative MS/BS and relative BFR than their WT controls, indicating a reduced response to loading. These data suggest that primary cilia in osteocytes not only serve as mechanosensors in bone but also play a key role in bone homeostasis, highlighting its potential as a therapeutic target in bone loss prevention.

The decreased rBFR and rMAR after loading due to the disruption of the primary cilia is consistent with prior studies, showing evidence that the primary cilia in osteocytes are necessary for load-induced bone formation [27]. Several studies have been conducted to explore the role of primary cilia in osteocytes in mechanotransduction, including one with an osteoblast- and osteocyte-specific deletion of Kif3a (one of the proteins that forms IFT anterograde motor Kinesin-2). The researchers observed that their cKO mice exhibited a significant decrease in bone formation after loading compared to their controls [28, 29]. With their mice models, however, no distinction can be made on whether these changes are due to the cilia-deficient osteoblast or osteocyte. Furthermore, Kif3a has been described to have other non-ciliary functions, so we cannot attribute the results solely on its role within the primary cilium. By using Dmp1-Cre as our recombinase, we are producing osteocyte-specific recombination. Deletion of MKS5, a cilia-specific protein in osteocytes creating dysfunctional cilia, greatly impacted bone formation in response to loading, which accentuates the primary cilium’s involvement in bone mechanotransduction.

The results of this study suggest that the osteoblast primary cilium affects bone development and growth. Both male and female cKO mice were visibly smaller than their WT controls, having significantly lower body weights as well as shortened femur lengths at 8 weeks of age. Although these significant phenotypes were present, DXA scans showed no significant differences between cKO and WT mice in either BMD or BMC. High variability within our cKO groups as well as a low number of experiment subjects could account for the lack of significance in our BMD data. In trabecular bone, female cKO mice had a significant decrease in MAR and BFR when compared to their WT controls. Although the males showed the same pattern, the difference did not reach significance, most likely due to the low number of test animals and large variations within the WT group. The BV/TV and MS/BS values of cKO mice showed no significant differences from WT animals. Several plausible factors could have contributed to these results, including decreased osteoblastic activity in the cKO groups, decreased osteoclastic activity in cKO groups, and compensatory mechanisms in bone turnover minimizing the difference between test groups. In accordance with the results from the thin sections, when analyzing trabecular bone from micro-CT scans, female cKO mice had a significant increase in trabecular bone number; however, since total bone volume did not change, this could be attributed to a change in the microarchitectural configuration of the bone. To determine whether this suggestion is accurate, a bending test would need to be performed. Bending tests would provide parameters of bone quality, determining if the cKO bone is weaker than the WT as predicted. Another plausible explanation for these results is decreased osteoclast activity, which would account for the high number of trabecular bone. No difference was observed in osteoclast number in either male or female groups, but in order to determine osteoclast activity, pathway studies would need to be performed. Micro-CT analysis of the femur showed no significant differences in cortical bone; however, both male and female cKO groups had a decreased average of cortical bone volume, surface, and area compared to their WT controls. It is possible that with a larger number of test subjects these non-significant differences could become more prominent, reaching a level of statistical significance.

The observed phenotypes of size of the mice as well as the length of their femurs is consistent with that of similar studies, in which deletion of the primary cilia using Prx1-Cre and IFT88^LoxP/LoxP^ directing cilia deletion to early limb bud mesenchyme led to decreased length of long bones in mice. Their reasoning behind it varied from defects in endochondral bone formation, alterations in differentiation and/or alterations in proliferation of different cell types [30, 31]. Another study focusing on the deletion of the primary cilia by targeting IFT88, this time using Col2a-Cre, a marker of chondrocyte differentiation, also observed a significant decrease in body size between cKO and control mice [30, 32]. Previous research using the same Cre recombinase but affecting IFT20 as opposed to IFT88, also noticed significant bone phenotypes; however, they noticed a significant decrease in BMD of their KO mice compared to their controls [31]. Altogether, this study indicates that primary cilia in osteoblasts are required for bone growth. Since primary cilia are involved in the coordination of several signaling pathways within the skeletal system, its potential as a target organelle for the treatment of bone loss diseases should continue to be explored.

This study has highlighted the role of primary cilia in bone health. Deleting MKS5 and IFT88 from specific bone cells resulted in impaired loading-induced bone formation and impaired bone growth respectively. This demonstrates that primary cilia are important in both bone mechanotransduction and development. This study provides a basis and motivation for a more in-depth investigation of the topic.

In future studies, it would be interesting to evaluate the mice at different time points, from birth to adulthood, to determine whether these differences would be attenuated or diminished. Diaphonization of newborn mice might also shed some light on the effects of the deletion of the primary cilium in skeletal development and growth. In order to see when the most marked differences occur, DXA scans and X-Ray images of the mice should be taken at different time points. Effects of cilia knockout in the proliferation, differentiation, and communication of each bone cell type should also be investigated. Valuable information could be gained with this experimental model through mechanical testing of bone tissues and cell pathway studies focusing on early bone development and bone turnover. Understanding the mechanisms of the primary cilium in bone development, adaptation, and mechanotransduction can contribute to the development of therapeutics targeting the organelle and aiding in the treatment of bone loss diseases.

In conclusion, this study highlights a critical role of primary cilia in bone development and mechanotransduction, suggesting that the presence of primary cilia in osteoblasts play an important role in skeletal development, and primary cilia in osteocytes mediate mechanically induced bone formation.

## Funding

project was supported by NIH R21 AR074012 (JL).

## Acknowledgments

We would like to thank the Indiana Center for Musculoskeletal Health (ICMH) Histology Core for assisting with histological preparation. We also thank Staci Engle and Ruchi Bansal for their technical support.

## Notes

### Competing Interest Statement

The authors have declared no competing interest.

## References

1. Yuan, X., R.A. Serra, and S. Yang, Function and regulation of primary cilia and intraflagellar transport proteins in the skeleton. Ann N Y Acad Sci, 2015. 1335: p. 78–99.

2. Berbari, N.F., et al., The primary cilium as a complex signaling center. Curr Biol, 2009. 19(13): p. R526–35.

3. Hoey, D.A., M.E. Downs, and C.R. Jacobs, The mechanics of the primary cilium: an intricate structure with complex function. J Biomech, 2012. 45(1): p. 17–26.

4. Anderson, C.T., et al., Primary cilia: cellular sensors for the skeleton. Anat Rec (Hoboken), 2008. 291(9): p. 1074–8.

5. Huber, C. and V. Cormier-Daire, Ciliary disorder of the skeleton. American Journal of Medical Genetics Part C: Seminars in Medical Genetics, 2012. 160C(3): p. 165–174.

6. Christensen, S.T., et al., Sensory Cilia and Integration of Signal Transduction in Human Health and Disease. Traffic, 2007. 8(2): p. 97–109.

7. Youn, Y.H. and Y.G. Han, Primary Cilia in Brain Development and Diseases. Am J Pathol, 2018. 188(1): p. 11–22.

8. Moore, E.R. and C.R. Jacobs, The primary cilium as a signaling nexus for growth plate function and subsequent skeletal development. J Orthop Res, 2018. 36(2): p. 533–545.

9. Temiyasathit, S. and C.R. Jacobs, Osteocyte primary cilium and its role in bone mechanotransduction. Ann N Y Acad Sci, 2010. 1192: p. 422–8.

10. Kim, J.H., et al., Wnt signaling in bone formation and its therapeutic potential for bone diseases. Therapeutic Advances in Musculoskeletal Disease, 2013. 5(1): p. 13–31.

11. Bonewald, L.F. and G.R. Mundy, Role of transforming growth factor-beta in bone remodeling. Clin Orthop Relat Res, 1990(250): p. 261–76.

12. Katagiri, T. and T. Watabe, Bone Morphogenetic Proteins. Cold Spring Harbor Perspectives in Biology, 2016. 8(6): p. a021899.

13. Zanotti, S. and E. Canalis, Notch Signaling and the Skeleton. Endocrine Reviews, 2016. 37(3): p. 223–253.

14. Jensen, V.L., et al., Formation of the transition zone by Mks5/Rpgrip1L establishes a ciliary zone of exclusion (CIZE) that compartmentalises ciliary signalling proteins and controls PIP2 ciliary abundance. Embo j, 2015. 34(20): p. 2537–56.

15. Anvarian, Z., et al., Cellular signalling by primary cilia in development, organ function and disease. Nature Reviews Nephrology, 2019. 15(4): p. 199–219.

16. Veland, I.R., et al., Primary Cilia and Signaling Pathways in Mammalian Development, Health and Disease. Nephron Physiology, 2009. 111(3): p. p39–p53.

17. Corbit, K.C., et al., Kif3a constrains β-catenin-dependent Wnt signalling through dual ciliary and non-ciliary mechanisms. Nature Cell Biology, 2008. 10(1): p. 70–76.

18. Li, C., et al., MKS5 and CEP290 Dependent Assembly Pathway of the Ciliary Transition Zone. PLoS Biol, 2016. 14(3): p. e1002416.

19. Roberson, E.C., et al., TMEM231, mutated in orofaciodigital and Meckel syndromes, organizes the ciliary transition zone. J Cell Biol, 2015. 209(1): p. 129–42.

20. Wang, L., et al., Ciliary gene RPGRIP1L is required for hypothalamic arcuate neuron development. JCI Insight, 2019. 4(3).

21. Haycraft, C.J., et al., Intraflagellar transport is essential for endochondral bone formation. Development, 2007. 134(2): p. 307–316.

22. Bivi, N., et al., Cell autonomous requirement of connexin 43 for osteocyte survival: Consequences for endocortical resorption and periosteal bone formation. Journal of Bone and Mineral Research, 2012. 27(2): p. 374–389.

23. Stratigopoulos, G., et al., Hypomorphism of Fto and Rpgrip1l causes obesity in mice. Journal of Clinical Investigation, 2016. 126(5): p. 1897–1910.

24. Corry, K.A., et al., Stat3 in osteocytes mediates osteogenic response to loading. Bone Rep, 2019. 11: p. 100218.

25. Blazek, J.D., et al., Disruption of bone development and homeostasis by trisomy in Ts65Dn Down syndrome mice. Bone, 2011. 48(2): p. 275–80.

26. Creecy, A., C. Smith, and J.M. Wallace, Dietary supplements do not improve bone morphology or mechanical properties in young female C57BL/6 mice. Scientific Reports, 2022. 12(1).

27. Moore, E.R., J.C. Chen, and C.R. Jacobs, Prx1-Expressing Progenitor Primary Cilia Mediate Bone Formation in response to Mechanical Loading in Mice. Stem Cells Int, 2019. 2019: p. 3094154.

28. Nguyen, A.M. and C.R. Jacobs, Emerging role of primary cilia as mechanosensors in osteocytes. Bone, 2013. 54(2): p. 196–204.

29. Hoey, D.A., J.C. Chen, and C.R. Jacobs, The primary cilium as a novel extracellular sensor in bone. Front Endocrinol (Lausanne), 2012. 3: p. 75.

30. Serra, R., Role of intraflagellar transport and primary cilia in skeletal development. Anat Rec (Hoboken), 2008. 291(9): p. 1049–61.

31. Lim, J., et al., Primary cilia control cell alignment and patterning in bone development via ceramide-PKCzeta-beta-catenin signaling. Commun Biol, 2020. 3(1): p. 45.

32. Song, B., et al., Development of the post-natal growth plate requires intraflagellar transport proteins. Dev Biol, 2007. 305(1): p. 202–16.

